# Description of a New Telonemia Genus and Species with Novel Observations Providing Insights Into its Hidden Diversity

**DOI:** 10.1101/2025.05.28.656552

**Authors:** Helena Mostazo-Zapata, Alex Gàlvez-Morante, Cédric Berney, Xènia Maya-Figuerola, Cristiana Sigona, David López-Escardó, Elisabet L. Sà, Dolors Vaqué, Daniel J. Richter

**Affiliations:** Institut de Biologia Evolutiva (CSIC-Universitat Pompeu Fabra), Barcelona, Spain; Department of Marine Biology and Oceanography, Institute of Marine Sciences (ICM), CSIC, Barcelona, Spain

**Keywords:** Telonemia, protists, taxonomy, phylogeny, eukaryotic diversity

## Abstract

Telonemia is a fascinating and understudied group of microbial eukaryotes known to have a vast diversity that is still uncharacterized. In fact, although they are thought to be the closest relatives of the eukaryotic supergroup SAR (Stramenopiles, Alveolata and Rhizaria), their diversity and biology are largely unexplored: to date, there are only seven described species in three genera, although there are estimated to be hundreds more unknown lineages. Here, we describe the isolation and characterization of two new strains, including a new genus (*Hyaliora molinica* gen. et sp. nov.) and a new species (*Telonema blandense* sp. nov.), and the re-isolation of a previously characterized telonemid, *Telonema subtile*, accompanied by new behavioral observations. We present morphological measurements highlighting differences among the isolates and a phylogenetic tree incorporating their 18S rRNA gene sequences. Furthermore, key aspects of their cell biology and structure are highlighted to provide insights into the evolution of TSAR. Since they are relevant not only phylogenetically, but also play a crucial role in food webs with some very abundant representatives in aquatic ecosystems, the findings of this study provide a further sampling and culturing of Telonemia to increase the knowledge of the hidden diversity and evolution of this mysterious group.

## Introduction

Telonemia, commonly referred to as ‘telonemids’, is a poorly characterized eukaryotic group that holds an informative phylogenetic position, as it has been recently identified as the sister to the supergroup SAR (Stramenopiles, Alveolata and Rhizaria), which may comprise up to half of eukaryote species diversity (Grattepanche et al., 2018). Together, telonemids and SAR form the TSAR group (Strassert et al., 2019). Telonemids represent an opportunity to study the origin and evolution of their morphological synapomorphies and cellular innovations. Study of telonemids also provides insights into eukaryotic diversification and helps infer traits of the common ancestor within the TSAR group (Tikhonenkov et al., 2022).

Until now, there have been only seven characterized species of Telonemia, in three different genera, although they were initially described over a century ago. The first species, *Telonema subtile* (also known as *T. subtilis*, a homotypic synonym), was described in 1913 by Griessmann from the marine habitat (more specifically, from crude cultures of *Ulva* and red algae from Roscoff, France and Naples, Italy) as a relatively small (6-8 μm long), colorless, elliptical, rigid-bodied flagellate without a contractile vacuole and with no close relationship with other known flagellates (Griessmann, 1913). Another report on this species was published a few decades later, providing a further detailed morphological description with light microscopy, which incorrectly placed Telonemia within the now outdated family Cyathomonadidae (Cryptophyta, Chromista) due to its morphological similarity (Hollande, 1950), followed by several further studies (Vørs, 1992, 1993). It was not until 2013 when a more detailed characterization was finally made, specifying its structural characteristics: cells contain mitochondria with tubular cristae, extrusomes, a telonemosome (a unique organelle considered to be a vesicle containing immature flagellar hairs), a nucleus, a multilayered cytoskeleton and adhesive fibers (Yabuki et al., 2013). It took almost a century from the first description to isolate and characterize a second species, from the surface marine waters of the inner Oslo fjord, described as *T. antarcticum* (Klaveness et al., 2005), although in 2015 it was renamed as *Lateronema antarctica* due its morphological and genetic differences with *Telonema*, resulting in a second genus (Cavalier-Smith et al., 2015). Later, in 2019, phylogenetic analyses revealed that Telonemia occupies a robust position in the tree of eukaryotes, as part of an expanded supergroup (TSAR) (Strassert et al., 2019). More recently, in 2022, a new study was published describing 3 new species of Telonema (*T. rivulare, T. papanine* and *T. tenere*) and two new species within the novel genus *Arpakorses: A. idiomastiga* and *A. versatilis* (Tikhonenkov et al., 2022). All together, these 3 genera represent a very low percentage of telonemid diversity, as there are estimated to be over a hundred marine and freshwater undescribed lineages of telonemids (Bråte et al., 2010).

Currently, our knowledge of telonemids comes from previous studies that indicate that they are phagotrophic heterotrophic biflagellate protists of pyriform shape with flagella emerging on opposite sides of a short protruding antapical rostrum or proboscis (Klaveness et al., 2005) (Shalchian-Tabrizi et al., 2006). We note that a later study (Tikhonenkov et al., 2022) defined the flagellar pole as the ‘apical’ end of the cell, but here the original description will be used. As described above, their unique structural characteristics are also a shared feature (Yabuki et al., 2013) (Klaveness et al., 2005). This intricate cytoskeleton among other structures and morphologies might mean telonemids have retained ancestral conditions lost in other sister groups (Klaveness et al., 2005).

Due to their wide distribution and abundance, telonemids most likely play important ecological functions in both freshwater (Boukheloua et al., 2024) and marine ecosystems (Shalchian-Tabrizi et al., 2006). They feed on a wide range of bacteria and pico-to nanophytoplankton (Bråte et al., 2010) and it seems they do not survive without their eukaryotic prey (Tikhonenkov et al., 2022). This feeding behavior positions them as important consumers within the microbial food web, contributing to nutrient cycling and energy flow in marine ecosystems.

In this study, three new strains of telonemids have been isolated. These include a proposed new genus and a new species from Mediterranean marine samples. In addition, *Telonema subtile* has been reisolated from Antarctic marine samples. A detailed description of their morphology and behaviors is presented together with statistically supported morphological differences between these three strains, accompanied by a phylogeny built with available 18S ribosomal RNA gene sequences of telonemids.

## Materials and Methods

### Isolation and culture maintenance

Cultures of *Hyaliora molinica* gen. et sp. nov. strain BEAP0326 and *Telonema blandense* sp. nov. strain BEAP0082 were isolated by diluting an initial sample from subsurface water sampled at the Blanes Bay Microbial Observatory (BBMO), a station located in the NW Mediterranean Sea about 1 km offshore (41°40′N, 2°48′E) over a water column of 20 m depth in February, 2024. The isolates of *H. molinica* were kept at 17 ºC with 12 h light:dark cycles using RS medium in a ratio of 1 (Nutrient Media Component - P5) to 50 (Non-Nutrient Media Component - P4) (Sigona *et al*., in preparation; https://mediadive.dsmz.de/medium/P4 and https://mediadive.dsmz.de/medium/P5) and contained a mix of unidentified prokaryotes other eukaryotes. The cultures of *T. blandense* were stored at 17 ºC in darkness using RS medium 1:10 and included a mix of unidentified prokaryotes and other eukaryotes. *Telonema subtile* strain BEAP0314 was isolated from Antarctic surface waters (66º20’S, 67º30’W) sampled on February 22, 2023 over a water column of 5 m depth and was kept at 4 ºC in darkness using RS medium 1:100.

### 18S rRNA gene sequencing

A sample of 50 mL per culture was used for 18S cloning. The cells were resuspended in the pellet after being centrifuged for 20 minutes at 13000 x g. DNA was extracted using the DNeasy PowerSoil Pro kit (QIAGEN). The 18S rRNA genes were amplified by PCR using universal eukaryotic primers 42F-1747R (CTCAARGAYTAAGCCATGCA - CCTTCYGCAGGTTCACCTAC) for *T. blandense* and *H. molinica* and 82F-1732R (GAAACTGCGAATGGCTC - ACCTACGGAAACCTTGTTACG) for *T. subtile*.

Amplification products were purified using the NZYGelpure kit (NZYTech), cloned using the TOPO-TA Cloning kit (Invitrogen) and transformed into *E. coli* cells following LacZα-complementation. Positive clones were selected and amplified by PCR with vector-specific primers M13F-M13R (GTAAAACGACGGCCAGT - CAGGAAACAGCTATGAC). Sequencing was performed by Eurofins genomics using Sanger sequencing. The resulting sequences were base called using phred (Ewing et al., 1998) with the parameters ‘-trim_alt “” -trim_cutoff 0.01’ and assembled with phrap (De La Bastide & McCombie, 2007) with the parameter ‘-repeat_stringency 0.4’ and the consensus sequence was exported with consed (Gordon et al., 1998). Assembled sequences were compared with EukRibo version 1.0 (Berney et al., 2022) through BLAST (Camacho et al., 2009) so as to identify each sequenced isolate.

### Phylogenetic placements

To place the three new isolates in the phylogenetic tree of Telonemia, all of the available telonemid sequences in GenBank (*National Center for Biotechnology Information*, s. f.) were collected and added to the three new sequences together with sequences from two independent outgroups (Haptophyta and Provora). For some representatives, a complete 18S rRNA sequence was extracted from contigs in EukProt assembled transcriptomes (Richter et al., 2022) and was used to replace the GenBank sequence: *Arpakorses idiomastiga*, comes from contig TRINITY_DN173_c0_g1_i12; the sequence of *Arpakorses versatilis* was constructed from contigs TRINITY_DN1176_c0_g1_i11, DN1176_c0_g1_i9 and DN3_c1_g1_i5; *Telonema subtile* Tel-1 (Tikhonenkov et al., 2022), from contig TRINITY_DN559_c0_g1_i16; and outgroup sequences *Nebulosomas marisrubri*, that was manually assembled from contigs TRINITY_DN14194_c3_g2_i1 and TRINITY_DN14194_c3_g1_i16; *Nibbleromonas quarantinus*, from contig TRINITY_DN8372_c10_g3_i1; *Ubysseya fretuma*, manually assembled from contigs TRINITY_DN9703_c1_g4_i2, TRINITY_DN9703_c1_g3_i6 and TRINITY_DN9703_c1_g7_i1. In three other cases, complete 18S sequences were extracted from whole genome shotgun sequences: *Gephyrocapsa pseudohuxleyi*, extracted from whole genome shotgun sequence AHAL01000301; *Chrysochromulina tobinii*, extracted from whole genome shotgun sequence JWZX01002122; and *Diacronema lutheri*, extracted from whole genome shotgun sequence JAAMXO010001095. The alignment was performed manually and ambiguously aligned positions were also manually trimmed following secondary structure models for the 18S (Wuyts et al., 2000). Next, a maximum-likelihood phylogeny was inferred with RAxMLGUI 2.0 (Edler et al., 2021) along with automatic thorough bootstrap replicates under the GTR+GAMMA substitution model. Finally, the tree was visualized and edited with iTOL (Letunic & Bork, 2021).

### Morphological analysis

Still and time-lapse images were collected using differential interference contrast (DIC) with a Zeiss Axiovert Inverted microscope equipped with a 63x oil-immersion lens. Subsequently, digital images were processed using Fiji software (Schneider et al., 2012) (Schindelin et al., 2012).

Measurements of cell length, cell width, food vacuole diameter and flagella length were taken in 20 cells of each strain and analyzed with R Studio (RStudio Team, 2020). For flagellar length classification, the longer visible flagellum was designated as the “long flagellum”, while the shorter was identified as the “short flagellum”. Packages used were tidyverse (Wickham et al., 2019), ggplot2 (Wickham et al., 2016) and patchwork (Pedersen, 2024) for data analysis and the generation of graphs. To identify significance in difference between the three isolates, ANOVA tests were run with a 0.05 significance level. Normality of residuals was checked using Shapiro-Wilks and homogeneity of variances with Bartlett test. In the cases in which ANOVA was significant, multiple comparisons were performed with the Tukey test to find interspecific differences.

For scanning electron microscopy (SEM), cells were fixed with 25% glutaraldehyde for 3 hours at room temperature and then seeded on membranes with pores sized 0.8 μm (WHA10417301, MERCK Chemicals and Life Science) via filtering. The samples were dehydrated in a graded ethanol series and critical-point dried with liquid carbon dioxide in a Leica EM CPD300 unit (Leica Microsystems, Austria). The dried filters were mounted on stubs with colloidal silver and then were sputter-coated with gold in a Q150R S (Quorum Technologies, Ltd.) and observed either with a Hitachi S3500N scanning electron microscope (Hitachi High Technologies Co., Ltd, Japan) at an accelerating voltage of 5 kV or with a Hitachi SU8600 field emission scanning electron microscope (Hitachi High Technologies Co., Ltd., Japan) in the Electron Microscopy Service of the Institute of Marine Science (ICM-CSIC), Barcelona.

## Results

### Phylogenetic relationships within Telonemia

We reconstructed a phylogenetic tree containing the 18S rRNA gene sequences of our three new isolates of telonemids, together with all currently available telonemid 18S sequences from described species and environmental surveys. Based on its 18S rRNA gene sequence, one of the three isolates corresponds to *T. subtile*, as it is 100% identical to *Telonema subtile* Tel-1 (Tikhonenkov et al., 2022). A second isolate is identified as a new species of *Telonema* (*Telonema blandense* sp. nov.), as it lies in a strongly supported group containing *T. rivulare, T. tenere* and *T. subtile*. Lastly, a new genus is proposed for the third isolate, *Hyaliora molinica* gen. et sp. nov. (Figure 1), as it represents a pair of 18S rRNA sequences that cluster together and do not branch with any other described genus of telonemids.

**Figure 1.**
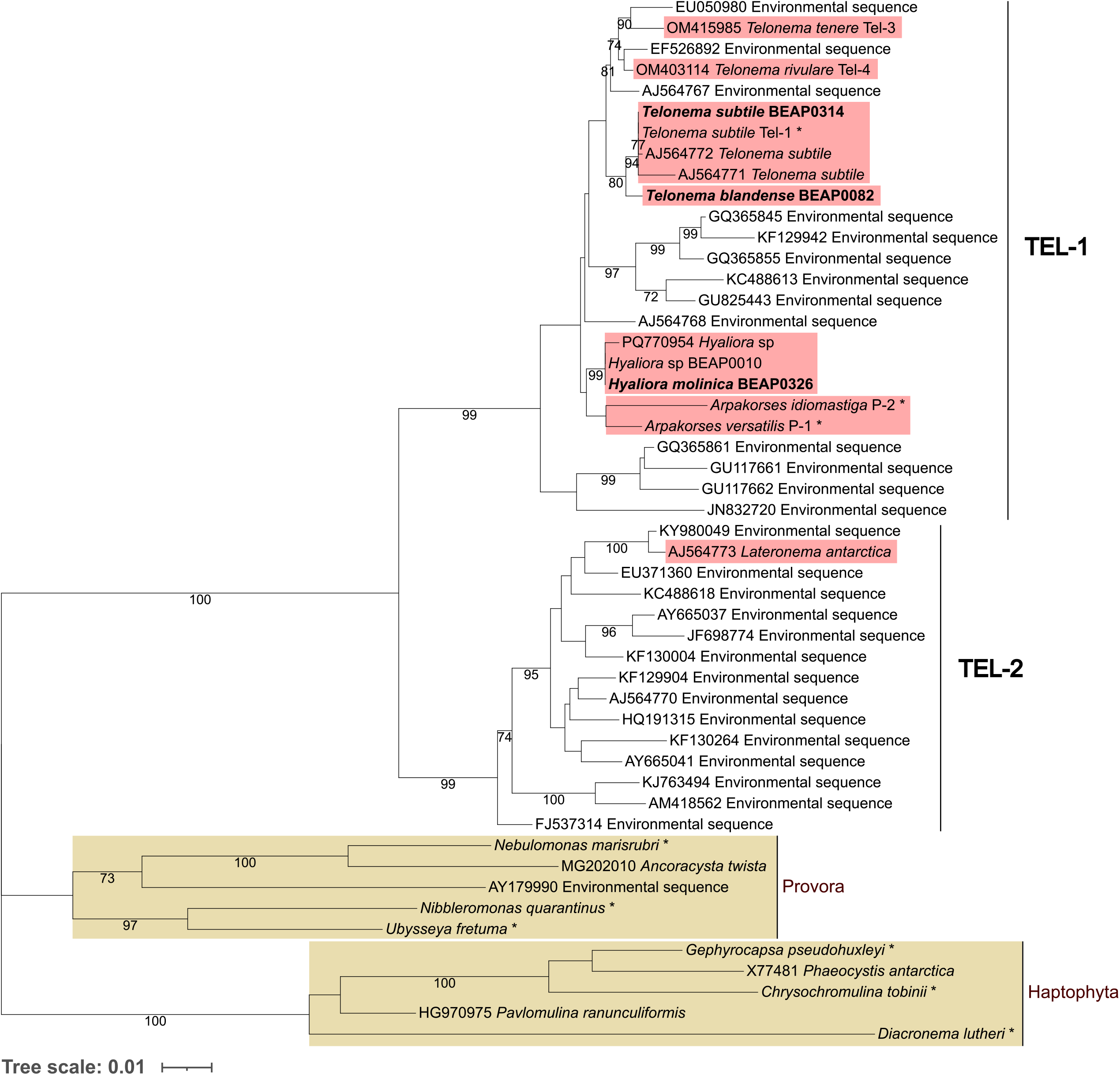
18S rRNA Telonemia maximum-likelihood phylogeny. Numbers at the nodes represent automatic thorough bootstrap (RAxML) support. Only values above 70 are shown. Names in bold represent the sequenced species in this work. Taxa in red boxes represent described species; whilst in brown, the two independent outgroups. Taxa with * represent complete sequences extracted from assembled transcriptomes or genomes (see Methods for details).

### Cell size and flagellar lengths: intraspecific variation and interspecific differences

In terms of cell size, *T. subtile* is significantly larger than *T. blandense* and *H. molinica* (length: p=10^−9^; width: p=10^−5^), which are not significantly different from one another (Figure 2a-b). As expected, food vacuole diameters are not significantly different among the three strains (p=0.3, Figure 2c), as they are likely to depend on food availability and prey size (Tikhonenkov et al., 2022).

**Figure 2.**
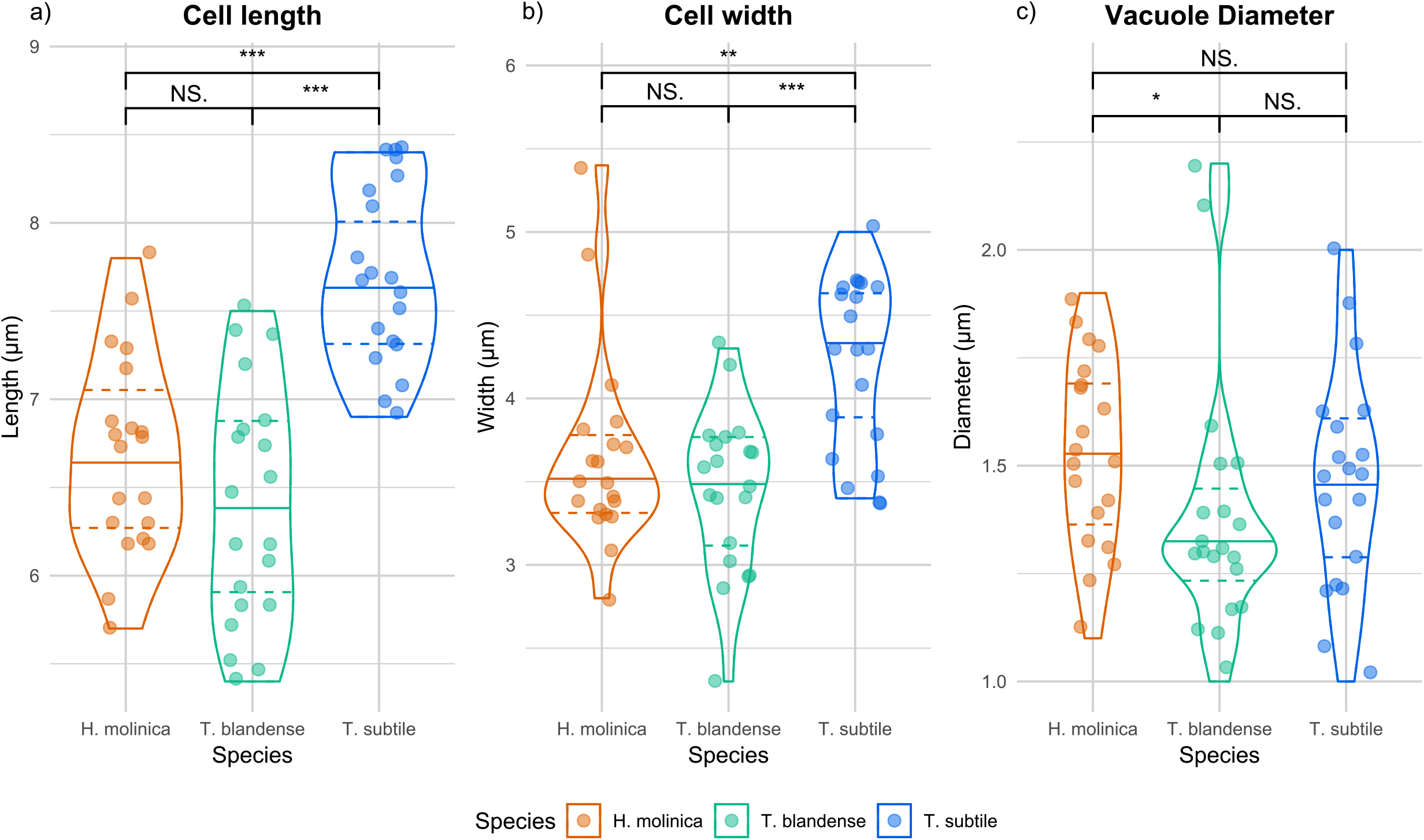
Distribution of cell length (a), cell width (b) and vacuole diameter (c) between the three isolates of this study. Data was collected from 20 cells with differential interference contrast (DIC) microscopy and measured using Fiji Software. * = p < 0.05; ** = p < 0.01; ** = p < 0.001; N.S. = Non-significant (p <!: 0.05).

In Figure 3 there is the distribution of values regarding the length of both long and short flagellum in each species (with the longer of the two flagella considered the “long flagellum”; see Methods). Multiple comparisons reveal *T. subtile* has a larger mean flagella length, followed by *T. blandense* and *H. molinica*. Moreover, it is clear that both *H. molinica* and *T. subtile* possess significantly uneven flagella (p=0.009). However, in *T. blandense*, both flagella would be equal, according to this data.

**Figure 3.**
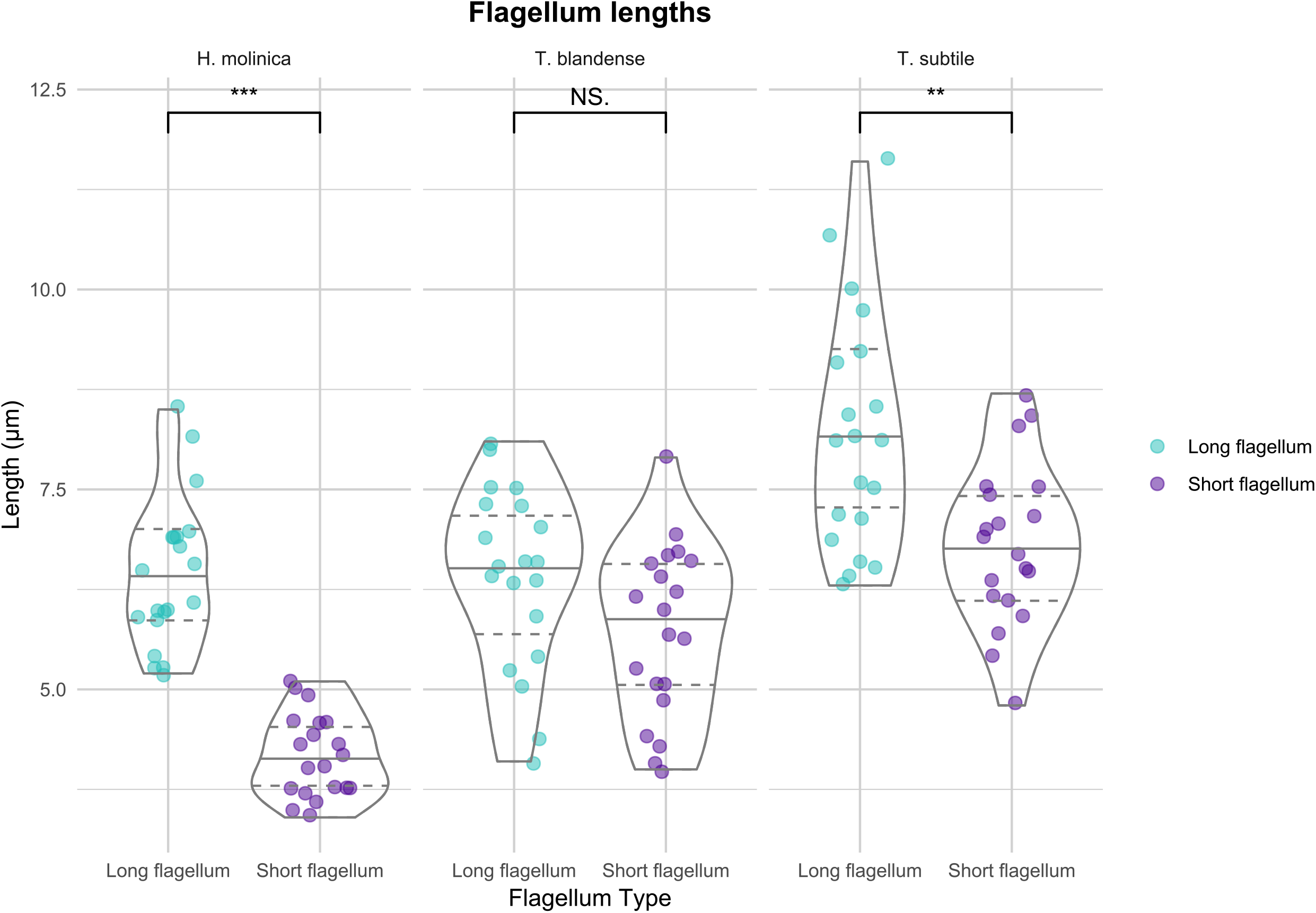
Distribution of flagella length between the three isolates of this study. Mean values are indicated in a blue point. Data was collected from 20 cells with differential interference contrast (DIC) microscopy and measured using Fiji Software. ** = p < 0.01; ** = p < 0.001; N.S. = Non-significant (p 0.05).

### External morphology, behavior and ecology of the isolates

We observed that all three telonemid strains in this study did not survive without another eukaryote present in the culture, presumably as prey. This has also been previously reported by (Tikhonenkov et al., 2022) in other isolates within Telonemia.

#### 1. *Hyaliora molinica* gen. et sp. nov

The cell body of *H. molinica* (Figures 4a-c and 5a-e) is drop-shaped with a rostrum containing a cytostome in the antapical part of the cell (Figure 5c,e), with a cell length of 5.7-7.8 μm and 2.8-5.4 μm in cell width. Two acronematic flagella emerge from the antapical part, with one of them (3.4-5.1 μm) significantly shorter than the other (5.3-8.5 μm) (Figure 3, Figure 4a). Mastigonemes were either missing or not detected in our images. In 7 out of 13 cells analyzed by scanning electron microscopy, we observed numerous protuberances in the base of the flagella (Figure 5c), although this observation may be an artifact of sample preparation. On the cell surface, a relief can be seen in all cells (Figure 5b, d, e) as well as a pit on one side of the cell (Figure 5e); the latter is a distinctive trait of telonemids (Tikhonenkov et al., 2022), (Shalchian-Tabrizi et al., 2006). A food vacuole is located in between the center and the posterior part of the cell (Figure 4a, c) and, in well-fed cells, it can be quite prominent, as reported by Klaveness et al. (2005) and Tikhonenkov et al. (2022) in other species of Telonemia. In those well-fed cells, the shape is rounder instead of pyriform. Cells swim twirling around their longitudinal axis with their flagella directed away from the direction of movement, although when sedentary, the flagella are wrapped around the cell body, with both in the same direction (Figure 4b). This behavior has been observed in species of the genus *Telonema*, but not in other genera (Tikhonenkov et al., 2022). Cell division is longitudinal, as described in all of the other characterized species of telonemids (Tikhonenkov et al., 2022).

**Figure 4.**
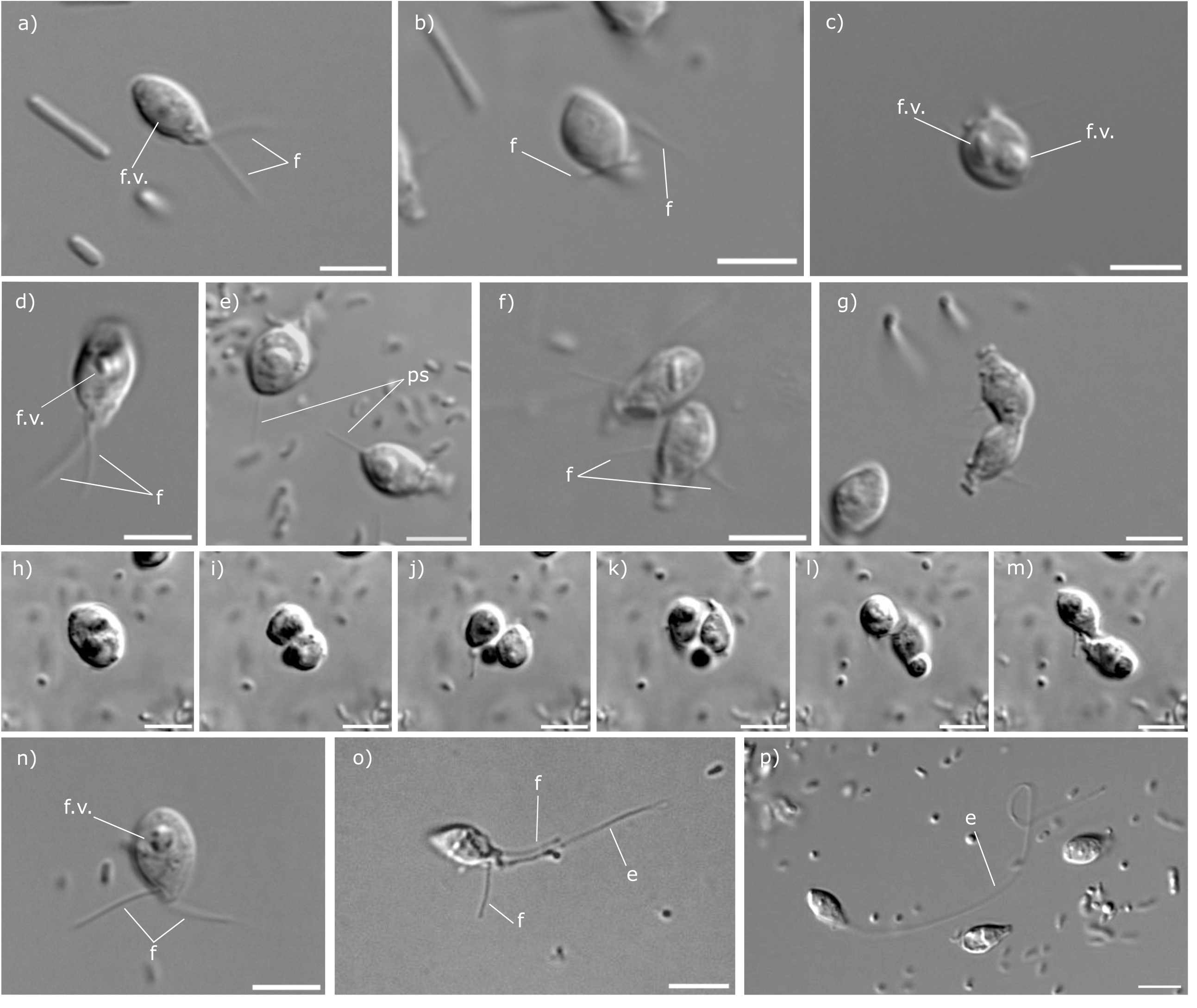
External morphology of the three telonemids isolated in this study with differential interference contrast (DIC) microscopy. *Hyaliora molinica*, free cell (a), attached to the surface (b) and in the first stages of division (c). *Telonema blandense*, free cell (d), two cells with pseudopodium (e), two cells attached to the surface (f), in the last stages of division (g) and sequence of possible vacuole reabsorption (h-m). *Telonema subfile*, free cell (m) and with elongating structure (n-o). Abbreviations: e - elongating structure; f-flagellum; f.v. - vacuole; ps - pseudopodium. Scale bar 5 µm.

**Figure 5.**
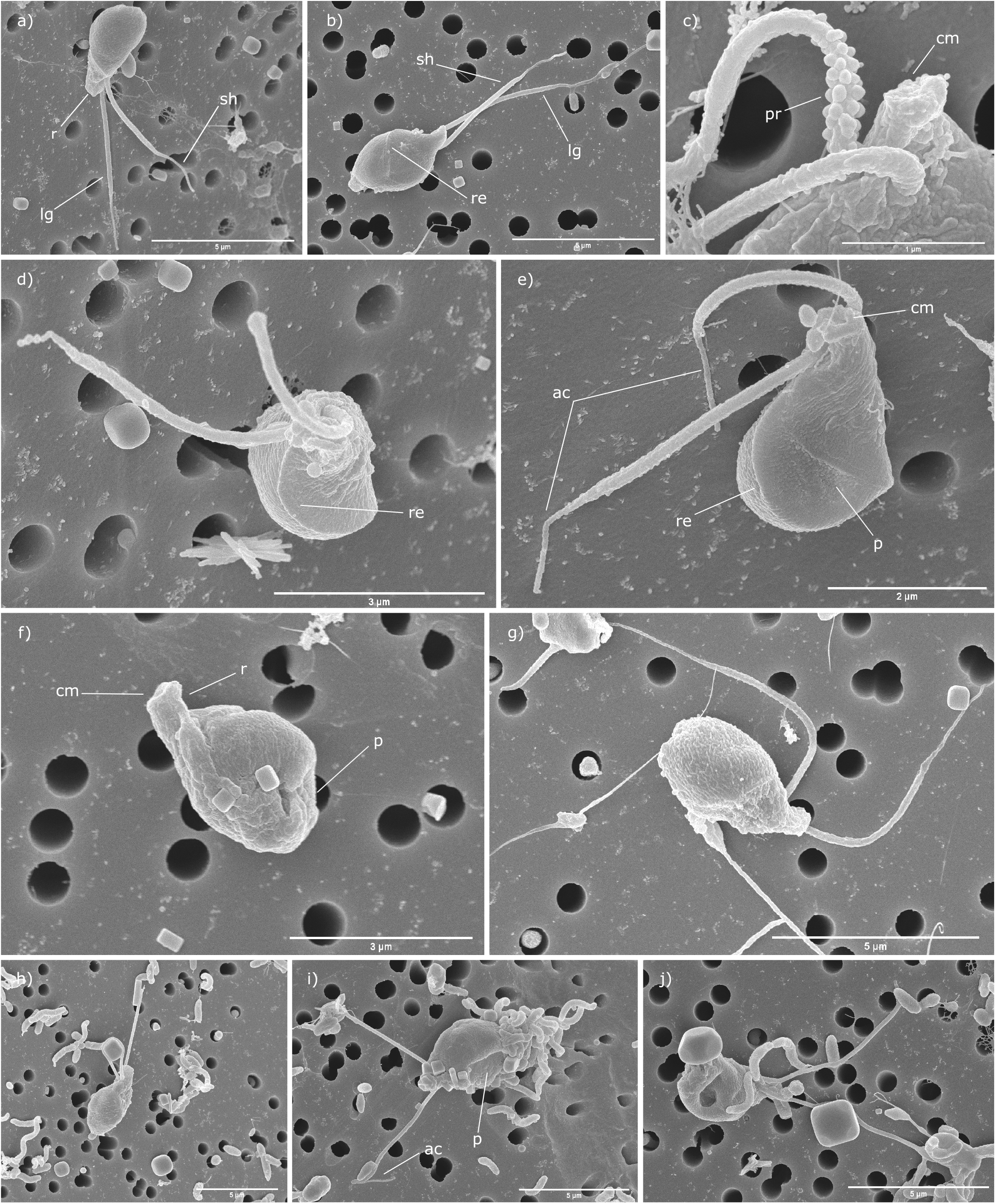
External morphology of the three telonemids isolated in this study with scanning electron microscopy (SEM). *Hya/iora molinica* (a-e), *Telonema blandense* (f-g) and *Telonema subfile* (h-j). Abbreviations: ac - acroneme; cm - cytostome; lg - long flagellum; p - pit; pr - protuberance; r - rostrum; re - relief; sh - short flagellum.

#### 2. *Telonema blandense* sp. nov

*Telonema blandense* sp. nov. is a proposed new species in the previously described genus *Telonema* (Figure 4d-m and Figure 5f,g) that possesses cell length (5.4-7.4 μm) and width dimensions (2.3-4.3 μm) similar to those of *H. molinica*. It has two emerging independent acronematic flagella that are not significantly different in length (3.9-7.9; 4.1-8.1 μm) (Figure 4d and 5g). No mastigonemes were detected. A cytostome in the rostrum can also be found in this species (Figure 5f) in addition to the pit in the surface. The food vacuole is consistently slightly closer to the posterior part of the cell in comparison to *H. molinica* (Figure 4d). A 0.7-5.6 μm long posterior pseudopodium was observed in sessile cells, where it appeared to provide substrate attachment (Figure 4e, Video 1 on [FigShare](https://doi.org/10.6084/m9.figshare.29066438)). They also swim spinning in their longitudinal axis with flagella backwards and wrap the flagella when not moving (Figure 4f). Cell division has been observed to be longitudinal (Figure 4g). Interestingly, in cells with a prominent vacuole we observed a new and unusual behavior: they seem to eject their vacuole to later reabsorb it, possibly as a way to free up space inside the cell during division (Figure 4h-m). These structures have also been seen being ejected from the cell and floating freely in the medium (Video 2 on [FigShare](https://doi.org/10.6084/m9.figshare.29066438)). To our knowledge, these observations have not previously been reported in any species of Telonemia.

#### 3. Telonema subtile

The body morphology and shape of *Telonema subtile* (Figure 4n-p and Figure 5h-j) is similar to all other characterized species within the genus, but with significantly bigger dimensions compared to *T. blandense* (and to *H. molinica*), with a cell length of 7.0-8.4 μm and width of 3.4-5.0 μm. It has two uneven acronematic flagella, also significantly larger than the other species in this study (5.7-8.7 and 6.3-11.6 μm). The food vacuole is consistently located in the posterior part of the cell, as in *T. blandense* (Figure 4n). In this species we have not observed wrapping of flagella when sedentary, consistent with previous studies (Tikhonenkov et al., 2022). However, a novel elongating structure in the antapical part independent from the flagella has been detected (Figure 4o,p, Video 3 on [FigShare](https://doi.org/10.6084/m9.figshare.29066438)). The structure was observed to be up to 71.2 µm in length, or 59.6 μm longer than the longer of the two flagella. Its function remains unknown (see Discussion).

## Discussion

The phylogenetic relevance of Telonemia has been previously highlighted (Strassert et al., 2019), despite the relative lack of information about its biology and diversity. In this study, we contributed to a better understanding of telonemids through the isolation and detailed analysis of a new genus, a new species of a previously characterized genus and the novel observations made in an already known species. These new isolates add to the 7 previously described species in the group.

Although there are morphological similarities among the strains described here, and indeed to previously described telonemids (Table 1), analysis of their 18S rRNA gene sequences demonstrate that they represent different genera and species. The three new strains we present possess all of the traits distinctive to the group, although there are some differences among them, such as a larger cell body size in *Telonema subtile* (Figure 2a). Also, flagella length is also distinctive and both *T. subtile* and *H. molinica* possess uneven flagella.

**Table 1.** Morphological comparison between described telonemids.

| Species | Isolation site and habitat | Cell length (µm) | Cell width (µm) | Vacuole diameter (µm) | Flagella length (µm) | Uneven flagella | Flagella wrap around the cell | Other features |
| --- | --- | --- | --- | --- | --- | --- | --- | --- |
| *Hyaliora molinica* | Blanes, Spain (41°40'N, 2°48'E), marine | 5.7-7.8 | 2.8-5.4 | 1.1-1.9 | 3.4-5.1; 5.3-8.5 | yes | yes | Cell surface relief |
| *Telonema blandense* | Blanes, Spain (41°40'N, 2°48'E), marine | 5.4-7.4 | 2.3-4.3 | 1.0-2.2 | 3.9-7.9; 4.1-8.1 | no | yes | Presence of posterior pseudopodium and vacuole reabsorption |
| *Telonema subtile* | Antarctica (69°19'S, 66°95'W), marine | 7.0-8.4 | 3.4-5.0 | 1.0-1.9 | 5.7-8.7; 6.3-11.6 | yes | no | Elongating structure |
| *Telonema subtile*<br>(Tikhonenkov et al., 2022) | Kara Sea (75.888 N, 89.508 E), marine | 7.0–9.4 | 3.5–6.1 | NA | 7.5–9.2 | no | no | Cells swim with sharp jerks and alternately change motion from rectilinear to rotation |
| *Telonema papanine*<br>(Tikhonenkov et al., 2022) | Franz Josef Land archipelago, Russia (80°37'46.800 N, 56°53'45.500 E), freshwater | 8.5-13.7 | 4.2-8.3 | NA | 7.4-10.9 | no | yes | Presence of pseudopodium to attach to substrate |
| *Telonema tenere*<br>(Tikhonenkov et al., 2022) | White Sea (66°29'58.78700 N, 35°9'42.20600 E), marine | 5.8-10.2 | 3.5-7.2 | NA | 3.5-8.5 | no | no | Cell body more elongated and slightly curved inwards, cannibalism observed |
| *Telonema rivulare*<br>(Tikhonenkov et al., 2022) | Sakhray River, Russia (44°10'26.700 N, 40°17'58.800 E), freshwater | 10-14 | 5.0-9.5 | NA | 4.0-8.5 | no | yes | Relatively short flagella |
| *Arpakorses versatilis*<br>(Tikhonenkov et al., 2022) | Kara Sea (75.888 N, 89.508 E), marine | 5.6-8.7 | 3.5-6.1 | NA | 7.7-15.5; 7.6-11.3 | yes | no | Cells attach to the substrate with one flagellum |
| *Arpakorses idiomastiga*<br>(Tikhonenkov et al., 2022) | Kara Sea (75.888 N, 89.508 E), marine | 5.6-8.7 | 3.5-6.1 | NA | 3.3-7.2 | no | no | Form cell clusters, free flagellum creates a flow of water |
| *Lateronema antarctica*<br>(Klaveness et al., 2005) | Inner Oslo fjord, Norway (59°25'00"N 10°34'00"E), marine | 8-16 | 6-12 | NA | Not mentioned but significantly longer than the cell | yes | NA | Presence of alveoli in the surface |

As the first species, *Telonema subtile*, was described over a century ago (Griessmann, 1913) without an associated 18S rRNA sequence, it is impossible to determine with certainty whether previously identified isolates assigned to this species or genus belong to the genetic lineage that is now known as *T. subtile* or even to the genus *Telonema*. Consequently, all past morphological, ultrastructural, and behavioral observations not linked to an 18S sequence provide possible phenotypes within Telonemia; however, they cannot be certainly linked to a specific taxon, which may explain the observed morphological variations found among them. The isolate identified in this study as *T. subtile* and the isolate of Tikhonenkov et al., 2022 (Table 1) share the same sequence and exhibit sufficient morphological similarity to the original description; therefore, are now classified as *Telonema subtile*, whilst the original organism described would have represented a more general characterization of Telonemia. Consistent with this idea, we note that the original description of this species by Griessmann indicated that the flagella were equal in length. The discrepancy with the observation of unequal flagellar lengths in our culture could therefore be explained by the fact that a subset of the cells do indeed have flagella of equal length, and by the possibility that Griessmann’s description may have been based on a sample of cells that did not represent the variation present in the species.

Ultrastructurally, all studied species in this work show the morphology that defines this general description of Telonemia: biflagellated tear-shaped unicellular organisms with a pit on the surface (Shalchian-Tabrizi et al., 2006). Nevertheless, some exclusive traits have been observed in these new isolates, which might contribute to the understanding of the evolution in TSAR supergroup and in the eukaryotic cell. For instance, in *Hyaliora molinica*, there is a clear relief on the surface of all cells. A notable feature in Telonemia is the presence of a very complex cytoskeleton organization in diverse layers. According to Cavalier-Smith et al. (2015), two major parts of the row of anterior microtubules in *T. subtile* resemble the two posterior centriolar roots in Excavata, which might indicate that this highly intricate cytoskeleton is an ancestral condition kept by Telonemia and reduced differentially in the rest of eukaryotes (Strassert et al., 2019). It can be hypothesized that this relief on the surface is actually the belt or “corset”, the point where the anterior and posterior layers of the cellular peripheric cytoskeleton join. This belt is made of electronically dense material and tubulin microtubules (Tikhonenkov et al., 2022). On the other hand, there is the supposition that this relief is provoked by the so-called “alveoli”, which are present in the *Lateronema* genus (Klaveness et al., 2005) and is a distinctive trait in the closely related group Alveolata (Cavalier-Smith, 2013), meaning that they could be homologous (Cavalier-Smith, 2003). These “alveoli” are flattened vesicles under the cortical membrane containing electron-dense material (its composition varies depending on the group, but in *L. antarcticum* it was observed that they are crystalline structures (Klaveness et al., 2005)), and externally, their relief can be observed, which resembles the relief in *H. molinica*. Their function would be to reinforce the cellular cortex (Cavalier-Smith et al., 2015). According to Takahashi et al. (2015), the common eukaryotic ancestor of the TSAR supergroup, Glaucophyta and Haptista would have these structures present in their cell body.

Focusing on the elongating structure observed in *Telonema subtile*, previous studies have noted the presence of a pseudopodium in *Telonema papanine*, a recently characterized species from the same genus, which emerges from the posterior part and serves to anchor the cell to the substrate in sedentary cells (Tikhonenkov et al., 2022). We observed a morphologically similar pseudopodium in *T. blandense*, suggesting that it may serve the same function. However, there are morphological differences between the pseudopodia of *T. papanine* and *T. blandense* in comparison to our observations in *T. subtile*. In contrast to *T. papanine* and *T. blandense*, the elongating structure is longer, slightly thicker and it undulates actively (Video 3 on [FigShare](https://doi.org/10.6084/m9.figshare.29066438)). This suggests that, although the elongating structure of *T. subtile* might be involved in surface attachment, we also speculate that it could be somehow involved in environmental or prey sensing. Gliding locomotion is unlikely to be one of the functions, as cells have been only seen locally twisting rather than actively moving. Such structures were not characterized in previous descriptions (Shalchian-Tabrizi et al., 2006; Yabuki et al., 2013; Tikhonenkov et al., 2022). Further characterization of this structure is needed to fully understand its composition, origin and function.

Regarding the vacuole reabsorption we observed in *Telonema blandense*, there are no previous reports on a similar process in any other described telonemid species. It may be an unusual behavior in well-fed cells with a prominent vacuole that are in the process of division, as a way of freeing up space inside the cytoplasm. Another hypothesis would be that this event consists of extrusive organelles, since their presence has been reported in ultrastructure analyses of *T. subtile* (Yabuki et al., 2013) and the *Arpakorses* genus (Tikhonenkov et al., 2022).

From the ecological point of view, there is evidence of a wide distribution of telonemids in marine ecosystems at high abundance (Klaveness et al., 2005). Previous studies describe telonemids as phagotrophic organisms that prey on a wide range of bacteria and pico-to nanophytoplankton (Bråte et al., 2010), which means they likely play a crucial role in ecosystems as a consumer of other microorganisms of similar or smaller size. Since none of the three species characterized in this study survived without other smaller eukaryotes present in the cultures, telonemids might be dependent on these prey-predator dynamics. Identical observations have been previously made in the genus *Arpakorses* (Tikhonenkov et al., 2022).

In conclusion, in this study we shed an additional ray of light on the hidden diversity of Telonemia, which remains a mysterious phylum that may have a relevant evolutionary importance in the history of the eukaryotic cell. Broader environmental sampling in both marine and freshwater ecosystems around the world is needed to establish more clonal cultures of uncharacterized new lineages and to continue to obtain new information of the already described telonemids. Also, further genomic and transcriptomic sequencing from clonal cultures and environmental samples is required to build a more solid and detailed phylogeny in order to fully comprehend the vast biodiversity in this enigmatic phylum.

## Taxonomic summary

### Eukaryota, Diaphoretickes, Telonemia

*Hyaliora* Mostazo-Zapata, Gàlvez-Morante and Richter, n. gen.

Description: Biflagellated tear-shaped protists with a rostral outgrowth and cytostome in the antapical part of the cell. They possess two uneven acronematic flagella. They are phagotrophic and have been observed to not survive without other eukaryotes.

18S rRNA sequencing and phylogenetic analysis reveal genetic distance from other published Telonemia sequences and low bootstrap clustering with *Arpakorses* sp., suggesting a new genus for these cultures.

Etymology: from Greek ὕαλος (“glass” or “transparent”) and derived from Latin -ora (“beauty” or “appearance”). Feminine.

Type species: *Hyaliora molinica*.

Zoobank registration: Described under the Zoological Code; Zoobank registration will be performed following peer review.

*Hyaliora molinica* Mostazo-Zapata, Gàlvez-Morante and Richter, n. sp.

Description: Cell average dimensions are 6.7 × 3.6 μm with significantly uneven acronematic flagella (3.4-5.1; 5.3-8.5 μm). The most common cell shape is tear-shaped, although it varies depending on feeding conditions. They swim rotating around their longitudinal axis with their flagella directed backwards, although in sedentary cells, flagella are seen wrapping the cell body. Cell division is longitudinal.

Etymology: Species name refers to the first author’s birth town (Molins de Rei).

Type locality: Subsurface water, Mediterranean Sea, Blanes Bay, Spain, 41°40’ N / 2°40’ E.

Type material: The name-bearing type (an hapantotype) is an SEM stub deposited in the Marine Biological Reference Collections (CBMR) at the Institut de Ciències del Mar (ICM-CSIC, Barcelona, Spain) under the catalog/accession number ICMCBMR000699 (Guerrero et al., 2023). This material also contains other eukaryotes and uncharacterized prokaryote species, which do not form part of the hapantotype.

Gene sequence: The SSU rRNA gene sequence of isolate BEAP0326 is deposited in Genbank as PV654195.

Cell culture: A culture containing *H. molinica* and other eukaryotes and bacterial species is publicly available and has been deposited in the Roscoff Culture Collection (November, 2024; currently awaiting assignment of RCC ID).

Zoobank registration: Described under the Zoological Code; Zoobank registration will be performed following peer review.

*Telonema blandense* Mostazo-Zapata, Gàlvez-Morante and Richter, n. sp.

Description: Pyriform or ovoid biflagellated protist of 6.4 × 3.4 μm with equal acronematic flagella (4.0-8.0; 4.1-8.1 μm). Cells wrap their flagella around the body when not moving and swim with flagella directing backwards. A long posterior pseudopodium can be found in sessile cells. Ejection of the food vacuole has been observed when longitudinally dividing.

Etymology: Species name refers to the sampling location “Blanes” (Lat.).

Type locality: Subsurface water, Mediterranean Sea, Blanes Bay, Spain, 41°40’ N / 2°40’ E.

Type material: The name-bearing type (an hapantotype) is an SEM stub deposited in the Marine Biological Reference Collections (CBMR) at the Institut de Ciències del Mar (ICM-CSIC, Barcelona, Spain) under the catalog/accession number ICMCBMR000700 (Guerrero et al., 2023). This material also contains other eukaryotes and uncharacterized prokaryote species, which do not form part of the hapantotype.

Gene sequence: The SSU rRNA gene sequence of isolate BEAP0082 is deposited in Genbank as PV654197.

Cell culture: A culture containing *T. blandense* and other eukaryotic and bacterial species is publicly available and has been deposited in the Roscoff Culture Collection (November, 2024; currently awaiting assignment of RCC ID).

Zoobank registration: Described under the Zoological Code; Zoobank registration will be performed following peer review.

This publication (work) will be registered with Zoobank following peer review.

## Data, code and materials availability

The 18S sequences of BEAP0010, BEAP0082, BEAP0314 and BEAP0326 have been deposited in GenBank with the accession codes PV654196, PV654197, PV654198, PV654195, respectively. Phylogenetic trees (and the data necessary to reproduce them), videos and SEM images have been deposited in FigShare at 10.6084/m9.figshare.29066438. Cell cultures containing BEAP0082, BEAP0314 and BEAP0326 (and mixed prey eukaryotes and bacteria) are publicly available and have been deposited in the Roscoff Culture Collection (November, 2024; currently awaiting assignment of RCC IDs). SEM stubs for BEAP0082, BEAP0314 and BEAP0326 have been deposited in the Marine Biological Reference Collections (CBMR) at the Institut de Ciències del Mar (ICM-CSIC, Barcelona, Spain) under the catalog/accession number ICMCBMR000700, ICMCBMR000698 and ICMCBMR000699, respectively.

## Acknowledgements

We thank Clara Cardelús, operating the Blanes Bay Microbial Observatory (BBMO), and Drs. Josep M. Gasol and Ramon Massana, for providing access to monthly water samples from Blanes Bay. We thank Helge Abildhauge Thomsen for valuable feedback on this manuscript. This work was supported by MCIN/AEI/10.13039/501100011033 and by the “European Union NextGenerationEU/PRTR” for electron microscopy, the European Research Council (ERC) under the European Union’s Horizon 2020 research and innovation programme (grant agreement No. 949745), the Proyecto de Generación de Conocimiento grants POLAR CHANGE (PID2019-110288RB-I00; PIs: M. Dall’Osto and R. Simó) and PID2023-152955NA-I00 funded by MICIU/AEI/10.13039/501100011033 and by ERDF/EU, and the Departament de Recerca i Universitats de la Generalitat de Catalunya (exp. 2021 SGR 00751).

